# Improved mammalian retromer cryo-EM structures reveal a new assembly interface

**DOI:** 10.1101/2022.03.04.482375

**Authors:** Amy K. Kendall, Mintu Chandra, Boyang Xie, William Wan, Lauren P. Jackson

**Affiliations:** Department of Biological Sciences, Vanderbilt University, Nashville, TN, USA; Center for Structural Biology, Vanderbilt University, Nashville, TN, USA; Department of Biochemistry, Vanderbilt University, Nashville, TN, USA

## Abstract

Retromer (VPS26/VPS35/VPS29 subunits) assembles with multiple sorting nexin (SNX) proteins on membranes to mediate endosomal recycling of transmembrane protein cargoes. Retromer has been implicated in other cellular events, including mitochondrial homeostasis, nutrient sensing, autophagy, and fission events. Mechanisms for mammalian retromer assembly remain undefined, and retromer engages multiple sorting nexin proteins to sort cargoes to different destinations. Published structures demonstrate mammalian retromer forms oligomers *in vitro*, but several structures were poorly resolved. We report here improved retromer oligomer structures using single particle cryo-electron microscopy (cryo-EM) by combining data collected from tilted specimens with multiple improvements in data processing, including using a three-dimensional (3D) starting model for improved automated particle picking in RELION. A retromer mutant (3KE retromer) that breaks VPS35-mediated interfaces was used to determine a structure of a new assembly interface formed by the VPS26A and VPS35 N-termini. The interface reveals how an N-terminal VPS26A arrestin saddle can link retromer chains by engaging a neighboring VPS35 N-terminus, on the opposite side from the well-characterized C-VPS26/N-VPS35 interaction observed within heterotrimers. The new interaction interface exhibits substantial buried surface area (∼7,000 Å^2^) and further suggests metazoan retromer may serve as an adaptable scaffold.

## Introduction

Eukaryotic cells contain multiple membrane-enclosed organelles, which dynamically communicate with each other through membrane trafficking pathways. Trafficking ensures exchange of lipid and protein cargoes between donor and acceptor compartments, and trafficking events are spatiotemporally regulated by multiple large multi-protein complexes. Retromer (VPS26/VPS35/VPS29 subunits) is one important complex that plays a well-established role in protein sorting at endosomes by maintaining dynamic localization of hundreds of transmembrane proteins that traverse the endocytic system. Mammalian retromer associates with the cytosolic face of endosomes, where it functions to recycle receptors, transporters, and adhesion molecules. Retromer is thought to mediate retrograde recycling to the *trans*-Golgi network (TGN) (*1–3*) or to the plasma membrane (*4–7*) through sorting into tubular-vesicular carriers (reviewed in (*8– 10*)). Retromer-dependent cargoes undergo selective trafficking to avoid sorting to late endosomes and subsequent degradation in lysosomes (*11*), which ensures cells maintain homeostasis of transmembrane cargoes at the plasma membrane and within the endolysosomal system. Retromer trafficking pathways become dysregulated in multiple ways, including aberrant protein processing (*12*) and sorting (*13, 14*). Disruptions to retromer trafficking in turn drive protein misfolding linked with neurological and neurodegenerative conditions, including Alzheimer’s disease (AD) (*15–17*), Parkinson’s disease (PD) (*18*), and Down’s syndrome (*19, 20*). Retromer pathways can also be hijacked by both bacterial and viral pathogens (*21, 22*).

Retromer is thus well established as an evolutionarily conserved heterotrimer that binds different Sorting Nexin (SNX) proteins to mediate cargo sorting from phosphatidylinositol-3-phosphate (PtdIns3*P*)-enriched endosomal membranes. Retromer binds multiple sorting nexin family members, including SNX-BARs, SNX3, and metazoan-specific SNX27 (reviewed recently in (*10, 23, 24*)). The retromer heterotrimer (formerly called Cargo Selective Complex, or CSC) has been implicated in direct cargo recognition in both mammalian cells (*2, 25*) and yeast (*1, 26*). More recently, SNX-BARs have been shown to bind cargo directly in mammalian cells (*27, 28*), and SNX3 promotes cargo recognition in both yeast (*29, 30*) and mammalian cells (*31*). Yeast SNX-BAR/retromer (*32*) and both yeast and mammalian SNX3/retromer (*33*) complexes have been shown to form tubules *in vitro*. SNXs may therefore be considered critical adaptors that promote membrane recruitment (*5, 34*); recognize cargo (*6, 27, 31, 35, 36*); and drive tubule formation (*34*).

From a structural viewpoint, multiple crystal (*31, 37–39*), single particle cryo-EM (*40*), and cryo-ET (*32, 33*) structures have revealed how retromer subunits fold in three-dimensional space; how subunits interact to form a stable and elongated heterotrimer; and how heterotrimers assemble *in vitro* to form higher order oligomers in vitreous ice (*40*) or assembled on membranes with sorting nexins (*32, 33*). Briefly, VPS35 adopts an elongated *α*-helical solenoid fold. VPS26 exists as three orthologues in mammals (VPS26A/VPS26B/VPS26C) and adopts a bi-lobal arrestin-like fold (*39, 41*). VPS26A and VPS26B bind the highly conserved VPS35 N-terminus (*39*), while VPS26C binds VPS35-like (VPS35L) in the retriever complex (*42*). VPS29 contains a metallophosphoesterase fold (*38, 43*) that serves as a scaffold for binding the α-helical solenoid of the VPS35 C-terminus (*31, 37, 40*). Cryo-ET reconstructions revealed how retromer assembles with either SNX-BAR or SNX3 on membranes. Retromer assembles on top of either SNX-BAR dimers or SNX3 to form high V-shaped arches mediated by interactions between VPS35 subunits. Back-to-back VPS26 dimers mediate interactions with SNX-BARs or SNX3. In solution, mammalian retromer forms multiple oligomers, including the retromer heterotrimer; dimers of trimers; a tetramer of trimers; and extended flat chains (*40*). The existence of multiple oligomeric assemblies suggests mammalian retromer may function as a flexible scaffold. Single particle and biochemical data revealed critical residues in the VPS35/VPS35 assembly interface, which was tested biochemically and in yeast (*40*) and subsequently reported in cryo-ET reconstructions (*33*). The cryo-ET and single particle reconstructions share several common themes. The mammalian VPS35/VPS35 dimer interface resembles yeast VPS35 dimers observed at the top of the V-shaped arches (*32, 33*). However, the curvature of the mammalian VPS35 interfaces observed in dimers and chains differs, which results in long and flat chains (*40*). Additionally, mammalian retromer appears to form a VPS26A-mediated “tip to tip” dimer (*40*) that differs from back-to-back VPS26 dimers observed in the presence of SNX3 (*33*) or the yeast SNX-BAR,Vps5 (*32*). The mammalian VPS26 “chain link” interface was poorly resolved because particles exhibit severely preferred orientation. These data suggest VPS26 may be capable of forming at least two different dimers, but the biological context of the “tip-to-tip” dimers remains unknown. Improved structural data for this putative interface is important so that structural models can be used to test function in cell culture or model organisms.

In this work, we report improved single particle cryo-EM reconstructions of multiple mammalian retromer oligomers, including the heterotrimer, dimers of trimers, and two VPS35/VPS35 sub-structures. These improved structural models arise from multiple improvements implemented during data collection and processing. Acquisition of tilted datasets provided critical missing views for some particles. For some structures, early rounds of automated particle picking using a three-dimensional (3D) starting model in RELION also provided additional views. Map sharpening followed by real space refinement in PHENIX resulted in improved heterotrimer reconstructions that allowed subunits to be assigned in maps more confidently. We further report a new structure of the N-VPS26A/N-VPS35 “chain link” interface using a retromer mutant (3KE mutant) that breaks VPS35 interfaces in chains (*40*). This new interface is formed by interactions between the VPS26A N-terminal arrestin fold with the N-terminus of VPS35 in a neighboring heterotrimer. The presence of this particle *in vitro* suggests retromer may be capable of forming a variety of assemblies, even in the absence of sorting nexin proteins.

## Results

### Retromer heterotrimer

To generate improved reconstructions, we combined images from previously published data (*40*) with a new third data set in which specimens were tilted to gain additional views (Figure S1; Table 1) (*44*). We obtained an improved heterotrimer reconstruction from 43,808 particles with an average resolution of 4.9 Å. Figure 1 formally compares this structure (panels A, C, E, G; (*44*)) with the previously published version (panels B, D, F, H; (*40*)). Critical missing views were acquired from the additional tilted data set (Figure 1E versus 1F). Map sharpening in PHENIX followed by several rounds of refinement and model building (see Methods) substantially improved the heterotrimer model (Figure 1A; 1G; Table 2). This structure demonstrates improved local resolution across the heterotrimer (Figure 1C, 1D). In the previous version (*40*), the N-VPS35/C-VPS26 interface was poorly resolved (Figure 1B, 1D) compared to the VPS35/VPS29 interface. In the new structure, the arrestin “saddle” fold of VPS26 is more clearly resolved (Figure 1A, 1G). Overall, improvements in both data collection and processing (see Discussion) played a key role in generating a higher quality reconstruction.

**Figure 1.**
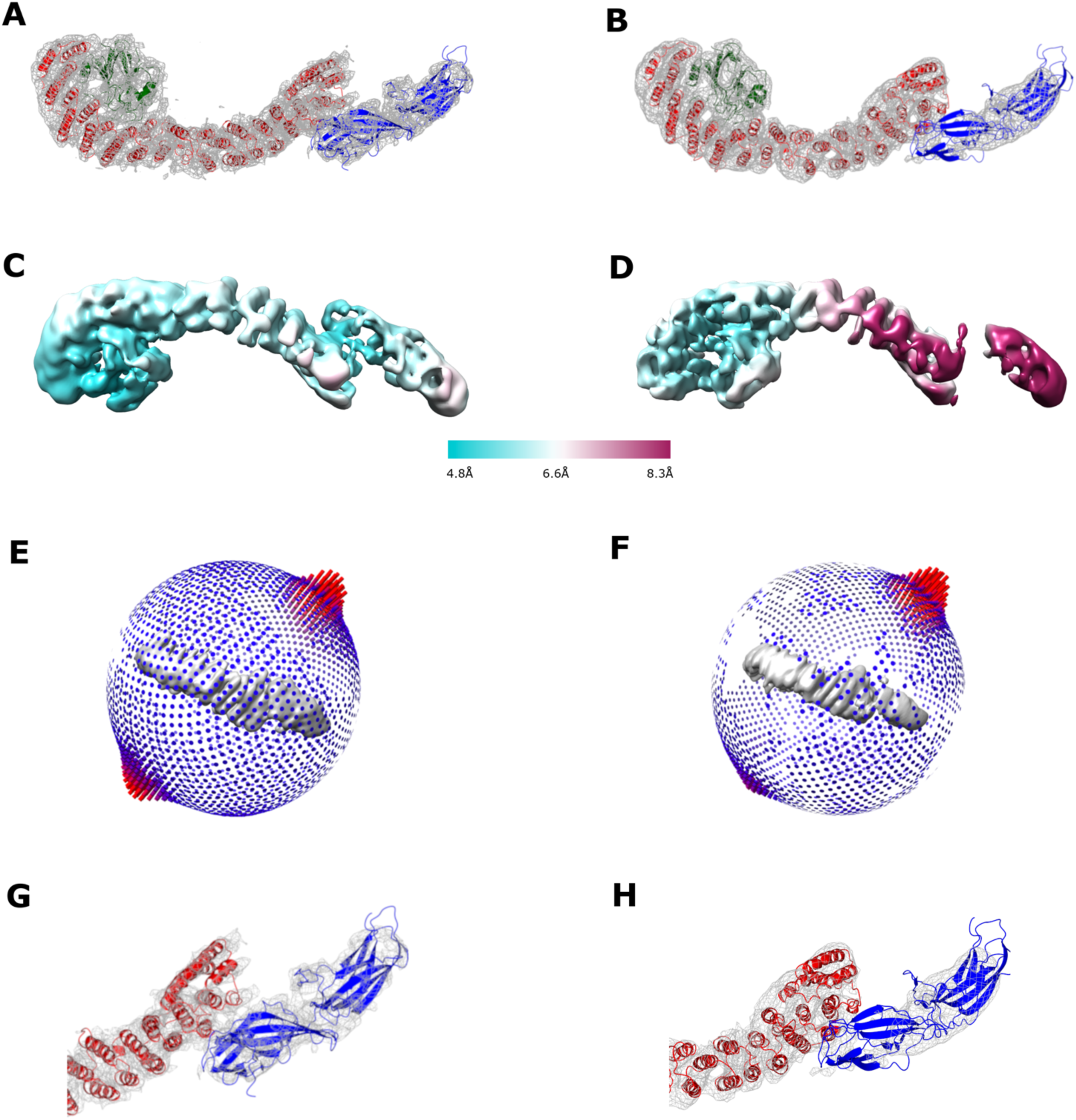
Single particle reconstructions of the mammalian retromer heterotrimer. (A, B) Comparison of new 4.9 Å resolution (A) and previously reported (B) reconstructions of retromer heterotrimer with fitted models. VPS29 is shown as green ribbons, VPS35 as red ribbons, and VPS26 as blue ribbons. (C, D) Local resolution comparisons across new (C) and reported (D) heterotrimer reconstructions. (E, F) Angular distribution in new (E) and reported (F) reconstructions. (G, H) Close-up views of N-VPS35/C-VPS26 interface in new (G) and reported (H) reconstructions. All Coulomb potential maps were generated using CCP4MG and are shown at 5σ contour level. Overall, acquisition of tilt data and improvements in data processing (details in text) resulted in an improved model.

### Retromer dimers of trimers

Retromer has been shown to form “dimers of heterotrimers” using both biophysical (*40, 45*) and structural (*40*) methods. In all reported conditions, formation of retromer dimers is mediated by the C-termini of VPS35 subunits. This dimer particle structure was very poorly resolved (18 Å average resolution) in previous studies (*40*). Others have reported improvements in single particle structure determination by using a 3D starting model (*46*) in early rounds of automated particle picking. We employed this strategy in RELION using a previously obtained data set (*40*) and a 3D starting model filtered to 20 Å to gain additional views of dimers in all possible orientations. We obtained an improved reconstruction (Figure 2A; Figure S2) from 288,247 particles with an average resolution of 7.0 Å. The presence of additional views substantially improved the structure (Figure 2E, 2F), although the N-terminal end of one heterotrimer remains poorly resolved and overall resolution remains limited compared to the heterotrimer (Figure 1).

**Figure 2.**
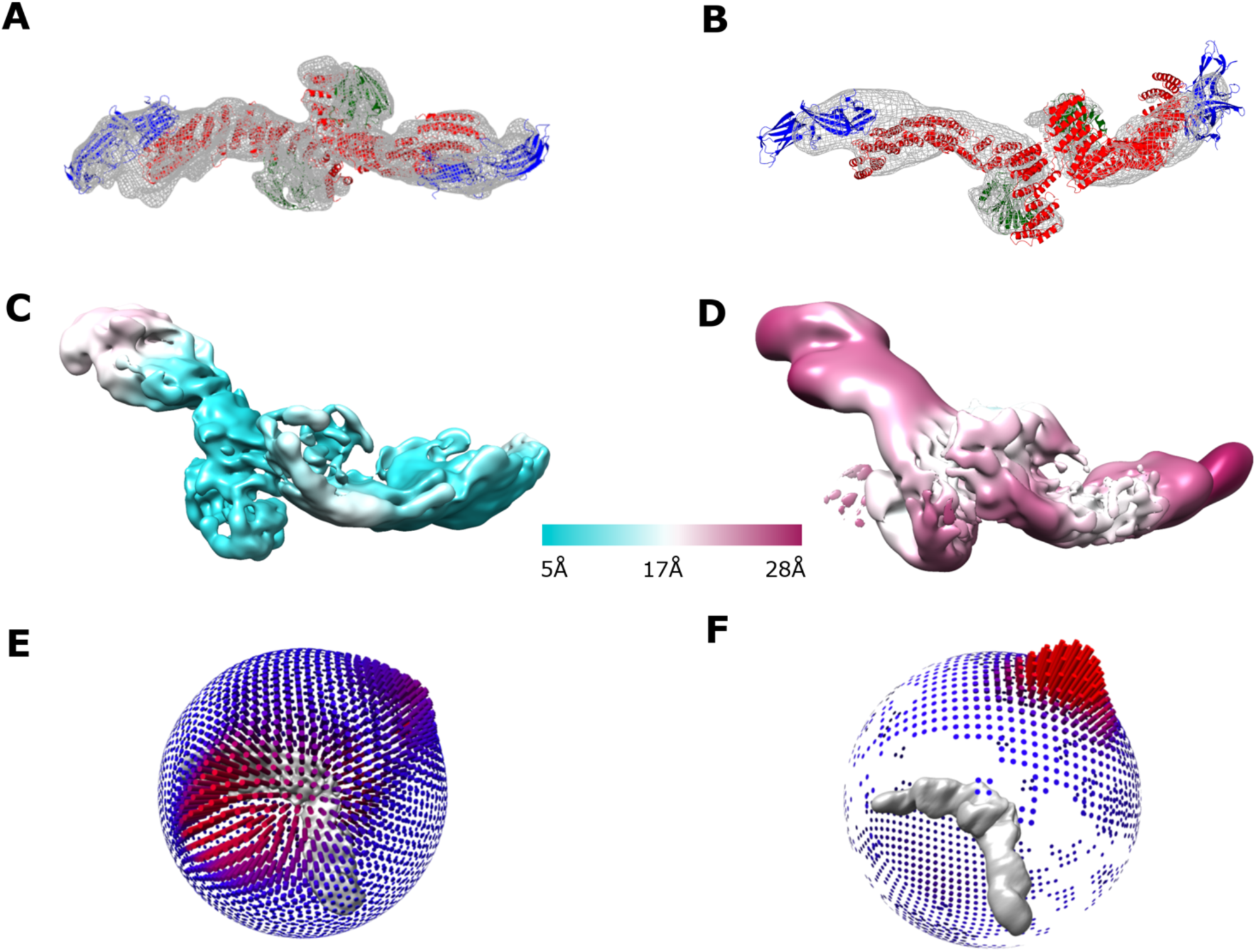
Single particle reconstructions of retromer dimers of trimers. (A, B) Comparison of new (A) and previously reported (B) reconstructions of retromer dimers of heterotrimers with fitted models. Coulomb potential maps generated using CCP4MG are shown at 6σ (A) and 4σ (B) contour levels, respectively. (C, D) Local resolution comparisons across new (C) and reported (D) reconstructions. (E, F) Angular distribution in new (E) and reported (F) reconstructions.

Fitting rigid body models into Coulomb potential maps can be ambiguous, especially at modest resolution. Therefore, we systematically analyzed map handedness in the improved dimer reconstructions to generate the best model given the data. Briefly, we undertook the following approach (*47*). Both the original and flipped maps were used to perform a global search in which 3,000 random orientations of the real space refined heterotrimer model (details in Methods) were fitted as rigid bodies. A list of potential fits and cross-correlation coefficients were generated for both maps (Figure S3A, D). Analyzing both maps with the top fitted model clearly demonstrated the map with correct handedness (compare S3B, C with S3E, F).

### VPS35/VPS35 sub-structures

Work from multiple labs has now demonstrated both yeast and mammalian retromer form VPS35-mediated dimer interfaces in solution (*40, 45*) and when assembled on membranes (*32, 33*). We re-visited two published sub-structures of VPS35/VPS35-mediated dimers (*40*) to generate improved reconstructions. The first sub-structure (Figure 3A) was determined from VPS35-mediated dimers observed in elongated retromer chains (*40*). The second (Figure 3B) was determined from retromer dimers after employing a 3D starting model in autopicking (previous section). The improved sub-structures revealed both interfaces are asymmetrical (C1 symmetry; Figure S1) and place specific electrostatic residues directly in the interface. These highly conserved residues (E615, D616, E617, K663, K701, K705) were identified and tested in our previous work (*40*), and improved sub-structures presented here now provide the highest resolution views of this critical VPS35-mediated assembly interface (see Discussion).

**Figure 3.**
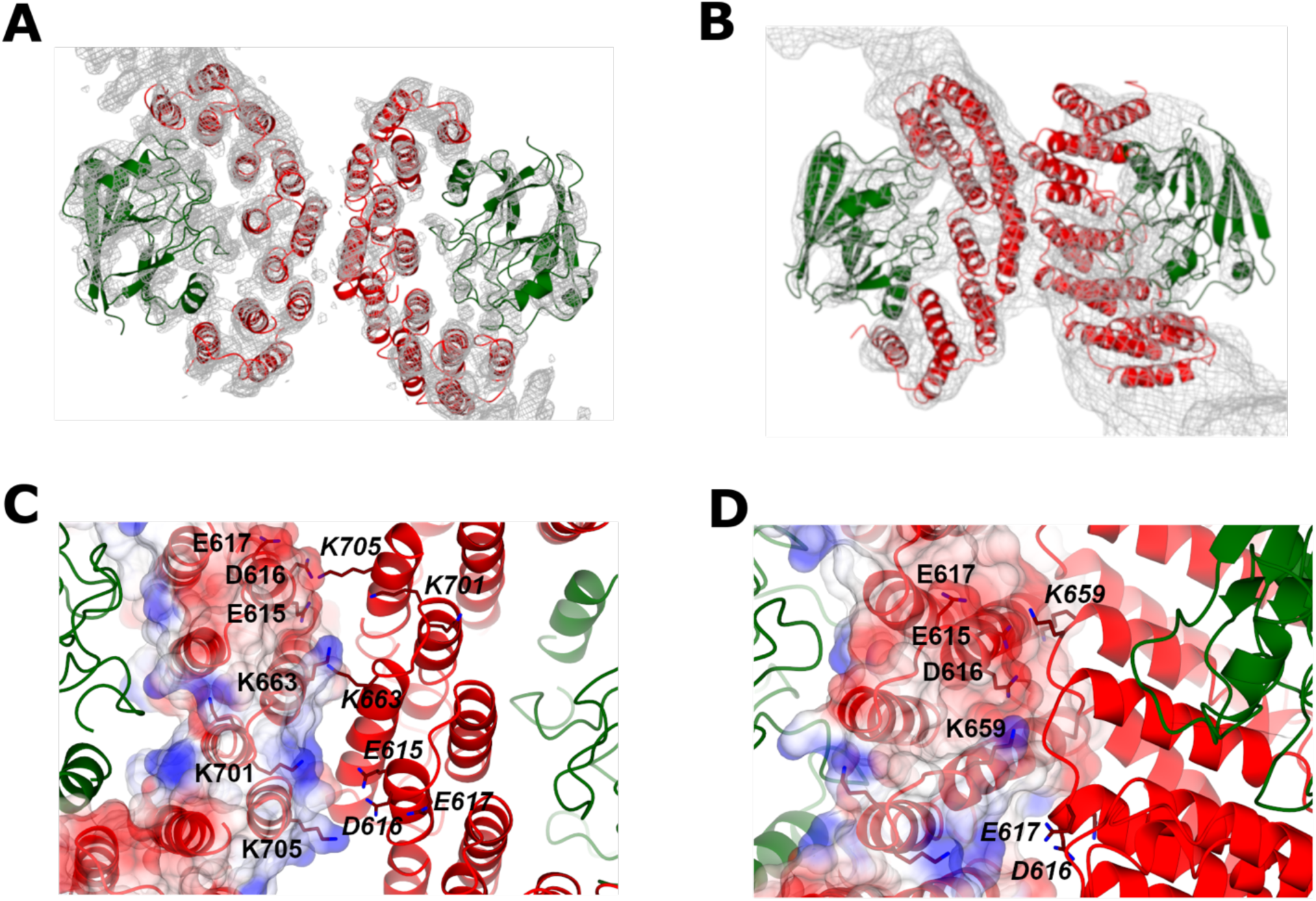
VPS35/VPS35 sub-structure reconstructions. (A) VPS35/VPS35 sub-structure interface determined from retromer chains is shown with fitted model. (B) VPS35/VPS35 sub-structure interface determined from retromer dimers of trimers (Figure 2) is shown with fitted model. Coulomb potential maps generated using CCP4MG are shown at 4σ (A) and 3σ (B) contour levels, respectively. (C, D) Close-up views of VPS35/VPS35 dimer interfaces observed in flat chains (C) and dimers of trimers (D). Improvements in data processing resulted in reconstructions with higher resolution; these structures further support the presence of specific electrostatic residues (E615, D616, E617, K663, K701, K705) mediating VPS35 dimer formation in asymmetric interfaces. Residues in the first dimer copy are labeled in black text, and residues in the second copy are labeled in black italic text.

### A new oligomeric interface mediated by VPS26A and VPS35

One major limitation of our previous study was an inability to resolve a new dimer interface found in chain links and mediated by the N-terminus of VPS26 subunits (chain interface II in (*40*)). These elongated chain particles exhibited extreme preferred orientation, so we used a published mutant that breaks the VPS35/VPS35 dimer interface to generate a particle more tractable for cryo-EM studies. This retromer “3KE” mutant (*40*) enriches the heterogeneous retromer protein sample for particles that form dimers only at the VPS26 end. These smaller 3KE retromer mutant particles resemble the “f-hole” found on the body of a violin (Figure 4). We obtained a tilted data set from 3KE mutant retromer in the presence of a small cyclic peptide called RT-L4 (*44*), which has been shown to stabilize retromer structure. The presence of this peptide further enriched our protein sample for 3KE mutant particles. We generated a reconstruction of 3KE mutant dimers (Figure 4A) from 40,957 particles using a 3D starting model (details in Methods) at an average resolution of 7.1 Å. One heterotrimer in this particle appears better resolved than the other (Figure 4C), which may reflect how the particle behaves at the air-water interface. The RT-L4 peptide is extremely hydrophobic, and 3KE particles with RT-L4 tend to exhibit preferred orientation by aligning along the plane of the interface (Figure 4E). To better resolve the interaction, we determined a sub-structure centered on the chain link from 19,386 particles at an average resolution of 6.7 Å. Although one copy of N-VPS35 remains more poorly resolved (Figure 4D), the sub-structure provides the first compelling view of this interface. The chain link is mediated by a new interaction between the N-terminal arrestin saddle of VPS26 in one heterotrimer with the N-terminus of VPS35 in a neighboring heterotrimer.

**Figure 4.**
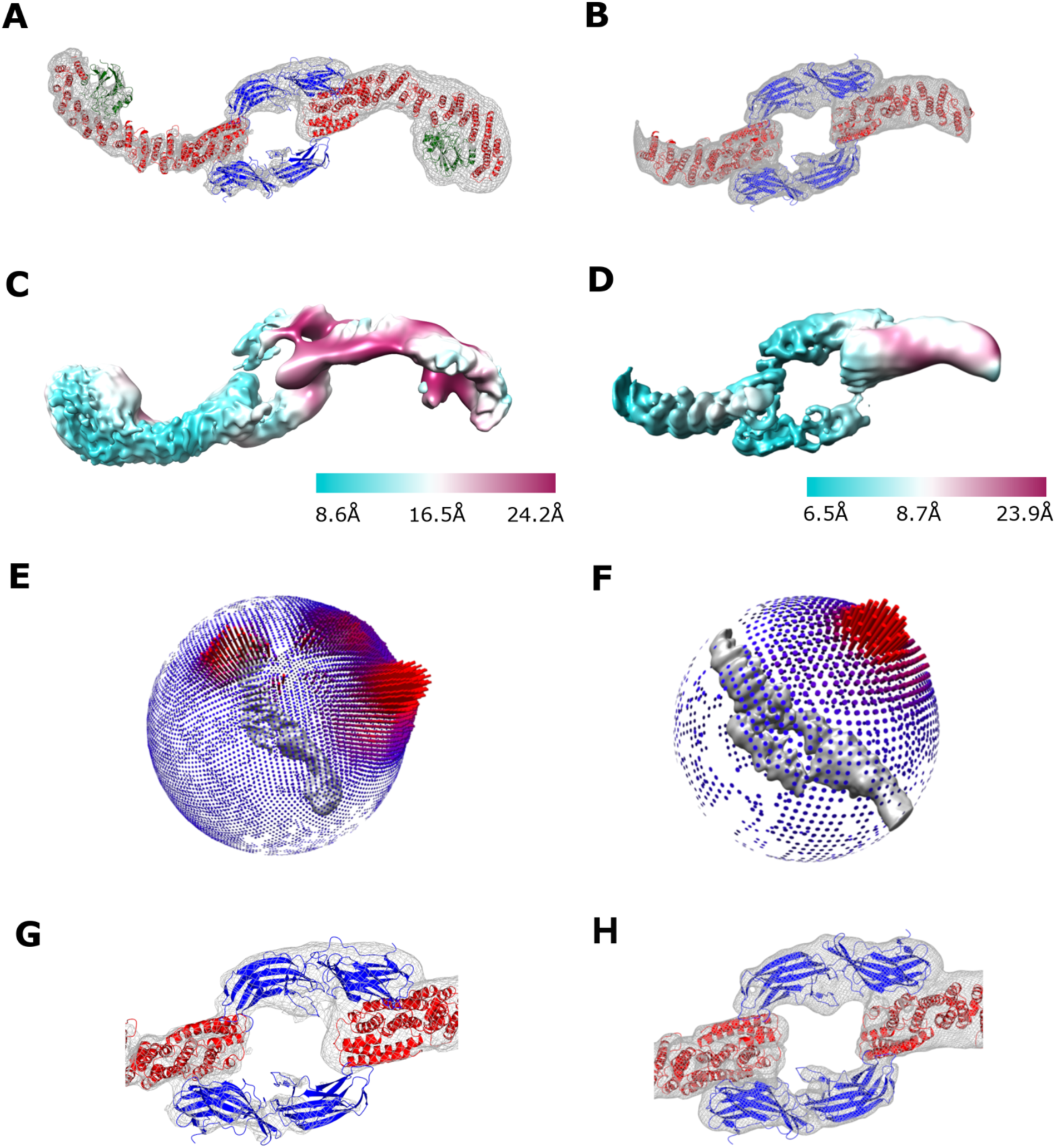
Single particle reconstruction of the retromer 3KE mutant reveals a new interface. (A, B) Reconstructions of retromer 3KE particle (A) and its sub-structure (B) centered on the “chain link” mediated by VPS26 and VPS35. Coulomb potential maps generated using CCP4MG are shown at 5σ (A) and 3σ (B) contour level. (C, D) Local resolution of 3KE mutant (C) and sub-structure. (E, F) Angular distribution of 3KE mutant (E) and sub-structure (F). (G, H) Close-up views of the newly observed VPS26/VPS35 interface, in which the N-terminal VPS26 arrestin saddle interacts with the N-terminus of a neighboring VPS35 subunit.

### Analysis of the N-VPS26 and N-VPS35 interface

To further characterize the chain link interface, buried surface area was calculated using PISA (*48*) (Figure 5). Within each retromer heterotrimer, interactions between N-VPS35 and C-VPS26 (Figure 5A; black & grey hatched circles) bury between 6,382 and 6,987 Å^2^ (Figure 5B). For comparison, the buried surface interface between N-VPS35 and C-VPS26 in the heterotrimer structure (Figure 1) is 6,178 Å^2^. The slightly larger buried surface area values for the equivalent interface in 3KE mutant particles is consistent with RT-L4 playing a stabilizing role in this interface (*44*). The two new interfaces are located on the opposite face of N-VPS35, where each N-VPS35 interacts with an N-VPS26 subunit from a neighboring molecule (Figure 5A; black and grey circles). Each new interface buries around 7,000 Å^2^ (Figure 5B). This analysis suggests the new interface buries approximately the same amount of surface as the well-characterized interaction between N-VPS35 and C-VPS26 within a retromer heterotrimer. The VPS35 N-terminal helix (residues 12-36) interacts specifically with two VPS26A *β*-strands (residues 48-56; 105-111) and their connecting loops (residues 56-63; 101-105) (Figure S6D, S6E). Like the C-terminal VPS35/VPS35 interfaces (previous section), this N-VPS26/N-VPS35 interface does not exhibit two-fold symmetry.

**Figure 5.**
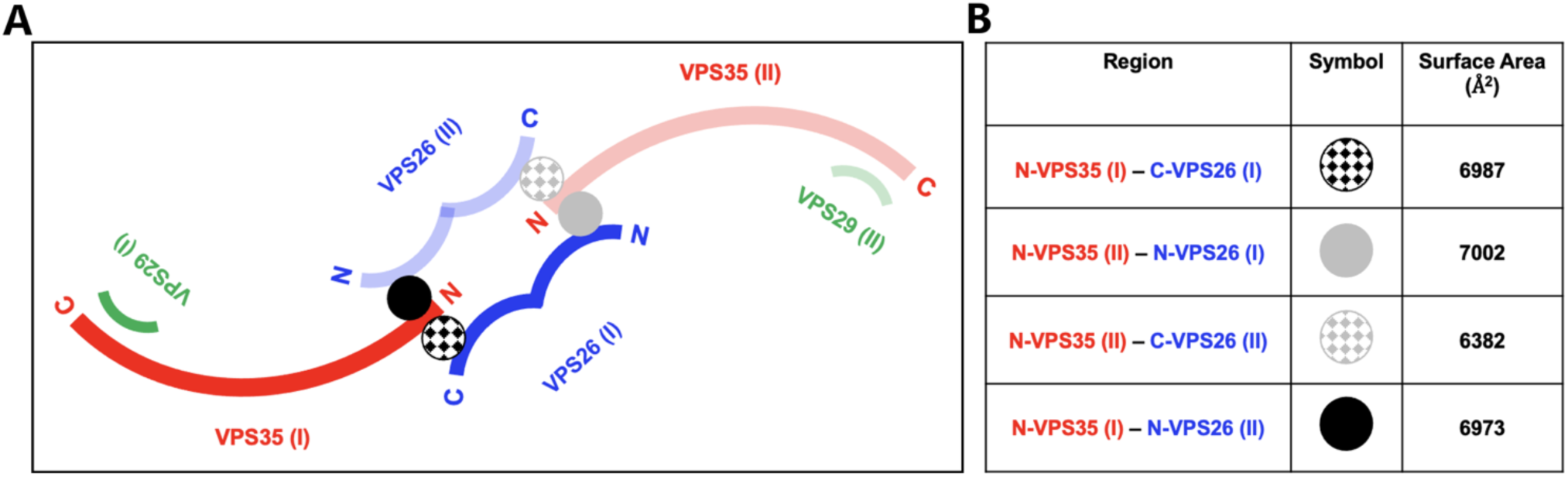
Analysis of retromer 3KE particle interfaces. (A) Schematic view of interactions observed in the retromer 3KE mutant particle. One heterotrimer is shown in dark colors (VPS29 in green, VPS35 in red, VPS26 in blue) and the second is shown in transparent colors. Within each heterotrimer, N-VPS35 binds C-VPS26A (black & grey hatched circles). Each “chain link” structure is held together by an interaction between a VPS35 N-terminus and the VPS26A N-terminal arrestin saddle in the neighboring heterotrimer (marked as black & grey circles). This interaction occurs on the opposite side from where the VPS35 N-terminus interacts with a VPS26A subunit within its own heterotrimer. (B) Summary of buried surface area analysis between VPS35 and VPS26A (calculated using PISA, full details in text).

## Discussion

### Improved data acquisition and image processing

Two major improvements allowed us to generate better reconstructions of retromer oligomers and to determine the structure of a new interface. First, data acquisition from tilted specimens retrieved multiple and important missing views for several structures, including retromer dimer and 3KE mutant particles. We previously proposed the VPS35 subunit could exhibit flexibility along the length of its *α*-helical solenoid, as judged by local resolution along the heterotrimer particle (*40*). However, the new reconstruction (Figure 1) instead suggests the poor local resolution at the N-terminus was instead driven by missing views. The decision to collect data from tilted specimens varies among practitioners, and our experience points to the importance of considering this as a data collection strategy. The second improvement arose from using a 3D starting model in early rounds of automated particle picking. Use of the 3D starting model helps programs identify and include poorly sampled views, and this was particularly effective in generating improved structures of retromer dimer and 3KE mutant particles.

### C-VPS35 dimer interfaces

The VPS35/VPS35 interface (Figure 3) we reported previously (*40*) includes several important conserved electrostatic residues observed at the top of retromer arches on membranes (*32, 33*). Both new sub-structures further support the placement of these residues in the interface. The improved structures also reveal the interface is not two-fold symmetrical, in agreement with reconstituted SNX3/retromer coats (*33*). We considered the possibility that VPS35-mediated dimers of heterotrimers observed in vitreous ice might represent V-shaped arches observed on membranes (*32, 33*). However, modelling arches into Coulomb potential maps after ascertaining correct handedness (Figure S6A) does not support this. Extended assemblies such as retromer dimers will likely be affected during sample blotting, so it is possible that arches observed on membranes have been flattened during the plunge freezing process by interactions at the air/water interface or by surface tension effects. Alternatively, dimers may represent a more loosely associated retromer structure that is further organized when it encounters relevant SNX proteins and cargoes on membranes.

### N-VPS26/N-VPS35 interface

The interface between the N-termini of VPS35 and VPS26A was previously observed in published structures of retromer chains (*40*) but is resolved more clearly in this work (Figure 4) by using a retromer mutant that breaks the long chains into smaller particles. The interface buries a substantial amount of surface area (nearly 7,000 Å^2^), suggesting it is a relatively stable particle *in vitro*. It is important to note this structure was determined in the presence of RT-L4 (*44*), a small molecule shown to stabilize the N-VPS35/C-VPS26 interface. The resolution of our structure is too low to locate or assign RT-L4, but in our hands, the presence of RT-L4 appeared to increase the number of 3KE retromer particles observed on grids. This suggests stabilizing the retromer heterotrimer in turn stabilized its ability to form the N-VPS26A/N-VPS35 interface.

The resolution of both the 3KE particle structure and its sub-structure (∼7 Å) are substantially lower than the C-VPS35 interface sub-structure determined from chains (∼4.5 Å). The limited resolution makes it difficult to accurately identify specific residues mediating the interface with high confidence since side chains cannot be assigned. Alpha-helices located in VPS35 interfaces (Figure 3A, 3B) can be more confidently fit, compared to the VPS26 *β*-sheets in the 7 Å resolution chain link sub-structure (Figure 4H). However, surface views of the interface suggest the two N-termini are complementary in overall shape (Figure S6D, E). We also note the VPS26 N-terminal arrestin subdomain is less conserved than the C-terminal subdomain (Figure S6F) (*39*), which may suggest retromer from different species cannot build this interface.

The biological relevance of the N-VPS26A/N-VPS35 interface remains unknown. It will be important to test these structural models by mutating multiple residues along the interface to ascertain whether disruption drives cargo sorting or other defects. In addition to endosomal sorting, retromer has been implicated in other important cellular processes, including mitochondrial derived vesicle formation (*49*) and regulation of mitochondrial membrane integrity (*50*), as well as nutrient sensing (*51, 52*) and autophagy (*53*). How retromer regulates mitochondrial morphology is unknown (*54*). Recent work (*55*) indicates a family of proteins called PROPPINs can compete with SNX proteins for retromer binding under certain conditions, and these PROPPIN-retromer interactions are linked to membrane fission events. Overall, the diversity of potential metazoan retromer functions suggests retromer may also have expanded its structural repertoire to assemble in different ways. This is not unprecedented, as other trafficking proteins have been shown to assemble in different ways. One well-studied example is clathrin, since clathrin on its own forms different lattices (*56*), and clathrin coated vesicles purified from brain exhibit a variety of geometries (*57*). It will be interesting to determine whether and how retromer may serve as a flexible scaffold to build different structures to mediate cellular events under a variety of conditions.

## Materials & Methods

### Reagents

Unless otherwise noted, all chemicals were purchased from Fisher (Waltham, MA, USA).

### Molecular biology and cloning

Both wild-type (*38, 39*) and 3KE mutant (*40*) retromer constructs have been published previously. Briefly, mouse VPS35 and VPS29 were placed in vector pGEX4T2 (GE Healthcare), while VPS26 was placed in in-house vector pMWKan. A two-stage quick-change mutagenesis protocol (*58*) was adapted to introduce three point mutations (K659E/K662E/K663E) into the VPS35 subunit to generate the retromer “3KE” electrostatic mutant that disrupts VPS35/VPS35 interface formation.

### Protein Expression and Purification

Recombinant retromer protein (wild-type or 3KE mutant) was expressed and purified from *E. coli* as previously described (*40*). Briefly, retromer plasmids were transformed into BL21(DE3) Rosetta2 pLysS cells (Millipore). Cells were grown to an OD_600_ between 0.8-1.0 and induced for 16-20 hours at 22°C with 0.4 mM IPTG. Cells were lysed by a disruptor (Constant Systems). The protein was purified in 10 mM Tris-HCl (pH 8.0), 200 mM NaCl, 2 mM *β*ME using Glutathione Sepharose (GE Healthcare). Protein was cleaved overnight using thrombin (Recothrom, The Medicines Company) at room temperature and batch eluted in buffer. Retromer was further purified by gel filtration on a Superdex S200 16/60 analytical column (GE Healthcare) into 10 mM Tris-HCl (pH 8.0), 200 mM NaCl, or dialyzed into 20 mM HEPES pH 8.2, 50 mM NaCl, 2 mM DTT prior to vitrification.

### Negative Stain Grid Preparation & Screening

For screening negatively stained retromer samples, 10 ml of retromer at concentrations between 5 and 10 μg/mL were applied to continuous carbon film on 400 square mesh copper EM grids (Electron Microscopy Sciences, Hatfield, PA) and washed twice with water. The grids were stained with 2% uranyl formate and 1% uranyl acetate and air dried overnight. The grids were screened on a ThermoFisher Morgagni microscope operating at 100 kV with an AMT 1Kx1K CCD camera (Center for Structural Biology Cryo-EM Facility, Vanderbilt University) to verify protein quality.

#### Cryo-EM sample preparation and data collection

*Grid preparation*. For cryo-electron microscopy, 2 μl retromer at a concentration between 80-100 μg/ml was applied to freshly glow discharged CF-2/2-2C C-Flat grids (Protochips, Morrisville, NC) or Quantifoil R 1.2/1.3 300 mesh copper grids (Quantifoil Micro Tools GmbH, Großlöbichau, Germany). Grids were vitrified in liquid ethane using either a ThermoFisher MarkIII or MarkIV Vitrobot, with blot times between 2-3.5 seconds and chamber conditions of 100% relative humidity and 8-20°C.

### Heterotrimer and flat substructure data collection

Micrographs from three separate data collections were used to generate the heterotrimer and flat substructure models. Each data collection is summarized below and full details are also provided in Table 1. RELION-2 (*59*) and RELION-3 (*60*) were used for all image processing unless otherwise indicated.

Data collection #1: 1480 micrographs were collected on a ThermoFisher Titan Krios microscope at the National Resource for Automated Molecular Microscopy (NRAMM). The microscope operated at 300 keV and was equipped with a Gatan BioQuantum energy filter with a slit width of 20eV, a spherical aberration (Cs) corrector, and a Gatan K2 Summit direct electron detector camera. The nominal magnification used during data collection was 105,000x, and the pixel size was 1.096 Å/pix. The total electron dose was 69.34e^-^/A^2^. Data collection was accomplished using Leginon (*61*). Images were motion corrected using MotionCor2 (*62*). The CTF of each micrograph was determined using Gctf (*63*); defocus values for the data varied between -0.7 and -2.6 μm.

Data collection #2: 1299 micrographs were collected on a ThermoFisher Titan Krios microscope at the National Resource for Automated Molecular Microscopy (NRAMM). The microscope operated at 300 keV and was equipped with a Gatan BioQuantum energy filter with a slit width of 20eV and a Gatan K2 Summit direct electron detector camera. The nominal magnification used during data collection was 105,000x, and the pixel size was 1.06 Å/pix. The total electron dose was 73.92e^-^/A^2^. Data collection was accomplished using Leginon. Images were motion corrected using MotionCor2, and micrographs from this data collection were rescaled to match the 1.096Å/pix pixel size from the first data collection using an NRAMM script written for MotionCor2. The CTF of each micrograph was determined using Gctf; defocus values for the data varied between -0.8 and -4.4 μm.

Data collection #3: 891 micrographs were collected on a ThermoFisher Titan Krios microscope at the National Resource for Automated Molecular Microscopy (NRAMM). The microscope operated at 300 keV and was equipped with a Gatan BioQuantum energy filter with a slit width of 20eV and a Gatan K2 Summit direct electron detector camera. The nominal magnification used during data collection was 105,000x, and the pixel size was 1.06 Å/pix. The total electron dose was 73.92e^-^/A^2^, and micrographs were collected at +/-15° tilts. Data collection was accomplished using Leginon. Images were motion corrected using MotionCor2, and micrographs from this data collection were rescaled to match the 1.096Å/pix pixel size from the first data collection using an NRAMM script written for MotionCor2. The CTF of each micrograph was determined using Gctf; defocus values for the data varied between -0.8 and -4.7μm.

### Dimer and dimer substructure data collection

1480 micrographs were collected on a ThermoFisher Titan Krios microscope at the National Resource for Automated Molecular Microscopy (NRAMM). The microscope operated at 300 keV and was equipped with a Gatan BioQuantum energy filter with a slit width of 20eV and a Gatan K2 Summit direct electron detector camera. The nominal magnification used during data collection was 105,000x, and the pixel size was 1.096 Å/pix. The total electron dose was 69.34e^-^/A^2^. Data collection was accomplished using Leginon. Images were motion corrected using MotionCor2. The CTF of each micrograph was determined using Gctf; defocus values for the data varied between -0.7 and -2.6μm. RELION-3 was used for all image processing unless otherwise indicated.

### Retromer 3KE mutant (f-hole) and sub-structure data collection

4791 micrographs were collected on a ThermoFisher Titan Krios G3i microscope in the Center for Structural Biology’s Cryo-EM Facility at Vanderbilt. The microscope was operated at 300 keV and equipped with a ThermoFisher Falcon3 direct electron detector camera. The nominal magnification used during data collection was 120,000x, and the pixel size was 0.6811 Å/pix. The total electron dose was 50 e^-^/A^2^, and micrographs were collected at +/-30° tilts. Data collection was accomplished using EPU (ThermoFisher). Images were motion corrected using MotionCor2. The CTF of each micrograph was determined using Gctf; defocus values for the data varied between -0.8 and -2.6 μm. RELION-3 was used for all image processing unless otherwise indicated.

#### CryoEM data processing

##### Heterotrimer

Data giving rise to our published retromer reconstruction (*40*) lacked tilted views; an additional data set (data collection #3) was collected to add tilted views and to improve the reconstruction for this study. Several thousand particles were manually selected from dataset #3 to perform initial 2D classification and produce templates for autopicking. Template-based autopicking in RELION identified 207,026 particles, which were subjected to initial 2D and 3D classification and refinement as well as CTF refinement. 250,500 particles from data collections #1 and #2 were imported to combine with data collection #3. Multiple rounds of 2D classification yielded 72,795 particles suitable to continue to 3D classification. Initial models for 3D classification were filtered to 60 Å resolution for use in these experiments. The particles underwent multiple rounds of CTF refinement and Bayesian polishing to produce a final set of 43,808 particles suitable for 3D refinement and postprocessing. The final masked heterotrimer model had a global resolution of 4.9 Å and a B-factor of -114 as determined in RELION.

##### VPS35/VPS35 sub-structure from retromer chains

Particles from the dataset containing all three data collections were windowed to a smaller box size that centered on the 35/35 interaction between two retromer molecules. Multiple rounds of 2D classification yielded 121,876 particles suitable to continue to 3D classification. Initial models for 3D classification were filtered to 60 Å resolution for use in these experiments. The particles underwent multiple rounds of CTF refinement and Bayesian polishing to produce a final set of 69,381 particles suitable for 3D refinement and postprocessing. The final masked flat substructure model had a global resolution of 4.5Å and a B-factor of -226 as determined in RELION.

##### Retromer dimers of trimers

Several thousand particles were manually selected to perform initial 2D and 3D classifications. Micrographs were then autopicked using a 20 Å low-pass filtered 3D model produced from initial rounds of manual picking; this template-based autopicking identified 533,231 particles. Multiple rounds of 2D classification yielded 509,447 particles suitable to continue to 3D classification. The same 3D model used for autopicking was filtered to 60 Å and used as an initial model for 3D classification. The particles underwent CTF refinement to produce a final set of 208,247 particles suitable for 3D refinement. The final masked dimer model had a global resolution of 7.0 Å, and no B-factor sharpening (postprocessing) was performed in RELION. We generated a z-flipped model of the dimer in Chimera (*64*). We then used the Chimera command “Fit in Map” to perform a global search testing 3000 random orientations of our real space refined heterotrimer model filtered to 7 Å, with random displacement of 5Å into each map. We generated a list of possible fits with cross-correlations for both the z-flipped and unflipped maps of the dimer.

##### Dimer substructure

The particles contained in the refined set from the full dimer were windowed to a smaller box size that centered on the VPS35/VPS35 interaction between two retromer molecules. Multiple rounds of 2D classification yielded 288,246 particles suitable to continue to 3D classification. A windowed model of the refined full dimer structure was filtered to 60 Å and used as an initial model for 3D classification. A set of 268,764 particles continued to 3D refinement, and the final masked dimer substructure model had a resolution of 6.5 Å; no B-factor sharpening (postprocessing) was performed in RELION.

##### Retromer 3KE mutant (f-hole)

Micrographs were autopicked using a 20 Å low-pass filtered 3D model that had been produced in early rounds of processing, and this template-based autopicking identified 275,633 particles. Multiple rounds of 2D classification yielded 154,553 particles suitable to continue to 3D classification. The same 3D model used for autopicking was filtered to 60 Å and used as an initial model for 3D classification. The particles underwent CTF refinement to produce a final set of 40,957 particles suitable for 3D refinement. The final masked model exhibited a resolution of 7.1 Å; no B-factor sharpening (postprocessing) was performed in RELION. To analyze map handedness, a z-flipped model of the retromer 3KE particle was generated in Chimera (Pettersen et al., 2004). We performed a global search testing 3,000 random orientations of the real space refined heterotrimer model (VPS26/VPS35/VPS29 as a single rigid body) filtered to 7 Å using the Chimera command “Fit in Map”, with random displacement of 5 Å into each map. This approach generated a list of potential fits with cross-correlations for both maps. Analyzing both maps with fitted rigid body models in Chimera clearly demonstrated the map with correct handedness.

##### Retromer 3KE (f-hole) substructure

The particles contained in the refined set from the full F-hole were windowed to a smaller box size that centered on the 35/26 tails of the two retromer molecules. Multiple rounds of 2D classification yielded 19,386 particles suitable to continue to 3D classification. A windowed model of the refined full f-hole structure was filtered to 60Å and used as an initial model for 3D classification. A set of 40,957 particles continued to 3D refinement, and the final masked f-hole substructure model had a resolution of 6.7Å; no B-factor sharpening (postprocessing) was performed in RELION.

##### Model Building & Docking

Models for VPS29 and the VPS35 C-terminus were obtained from PDB 2R17, while models for N-VPS35 and VPS26A were obtained from PDB 5F0J. We omitted a flexible unstructured loop (amino acids 470-482) that is absent in all crystal structures. Rigid-body docking and map visualization were performed in Chimera using the command “Fit in Map”.

##### Retromer heterotrimer map sharpening

We used the 5 Å resolution Coulomb potential map to refine an atomic model of retromer. The Coulomb potential map was further postprocessed in PHENIX by performing global or local B factor sharpening using the program “Auto-sharpening” with a resolution cut-off at 5 Å. A PDB model containing all three retromer subunits (VPS26 and VPS35 from PDB: 5F0J; VPS29 from PDB: 2R17)was docked into the sharpened map and traced manually in Coot. Subsequently, several rounds of real space refinement were performed in PHENIX (*65*). Overall statistics and geometry for the final model were analyzed using MolProbity (*66, 67*) and are summarized in Table 2. The overall root-mean-square deviation (RMSD) between C_α_ atoms in the refined versus starting models was 1.1 Å. However, the overall geometric parameters of C_α_ atoms are improved in the new model, suggesting the side chains likely occupy more favorable conformations in the new model. While overall improvements in the model appear relatively small, the overall improvement in the sharpened map is substantial, showing that with current tools a cryo-EM map of modest quality at 5 Å resolution can be used to generate a fairly reliable model (Figure 1).

##### Data visualization

All figures were generated using CCP4MG (*68*) or Chimera (*64*).

##### Data deposition

Coulomb potential maps were deposited in the EMDB with accession numbers EMD-24964, EMD-26342, and EMD-26341, corresponding to retromer heterotrimer; dimers; and 3KE mutant particles. Sub-structure maps were deposited for C-VPS35 dimers as EMD-26343 and EMD-26345, and for the 3KE mutant as EMD-26340. The heterotrimer data set was first reported very briefly in (*44*), but its refinement and analysis are reported here. Coordinates for this updated retromer heterotrimer were deposited in the Protein Data Bank as XXXX.

## Acknowledgements

The authors thank Kevin Chen and Brett Collins for generously providing the RT-L4 peptide used in 3KE retromer sample preparation and data collection. AKK, MC, BX, and LPJ are supported by NIH R35GM119525. WW is supported by NIH DP2GM146321. Screening was conducted at the Center for Structural Biology Cryo-EM Facility at Vanderbilt University. Data collection was performed at the National Center for CryoEM Access and Training (NCCAT) and the Simons Electron Microscopy Center located at the New York Structural Biology Center, supported by the NIH Common Fund Transformative High Resolution Cryo-Electron Microscopy program (U24 GM129539) and by grants from the Simons Foundation (SF349247) and NY State Assembly. The authors declare that they have no conflicts of interest with the contents of this article.

---

**Figure S1.**
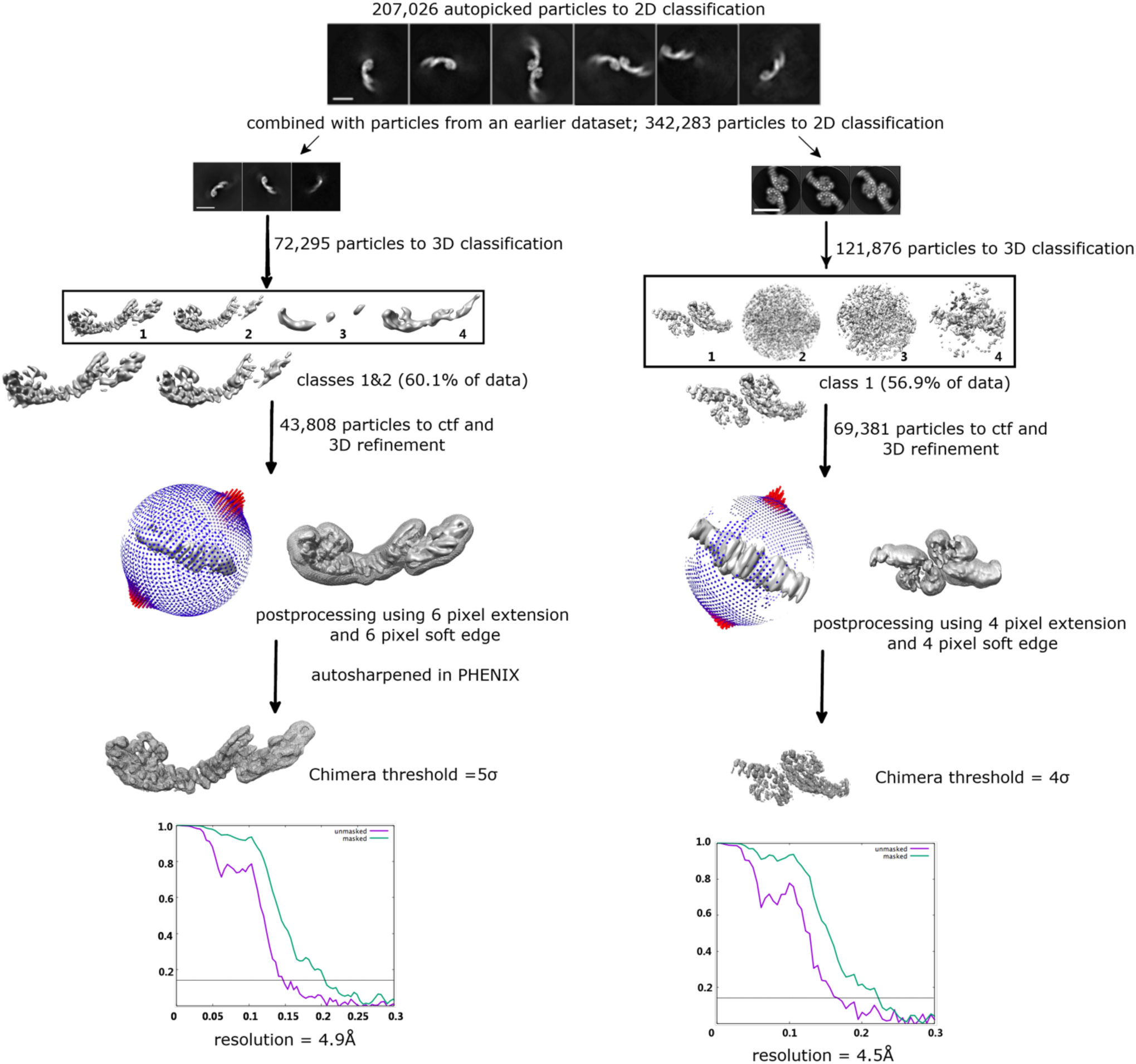
CryoEM image and data processing work flow for updated retromer heterotrimer and VPS35/VPS35 sub-structure. Particles were auto-picked from three combined datasets using a 3D starting model (details in Methods). Particles were separated into 2D classes based on biochemical species (heterotrimer, flat chains). The left branch shows data processing pipeline for heterotrimers, and the right branch shows processing for the VPS35/VPS35 sub-structure reconstructed from flat chains. 3D reconstructions were generated for each species. Fourier Shell Correlation (FSC) plots showing masked and unmasked resolution estimates from RELION are shown for each structure or sub-structure; the grey line marks the 0.143 cut-off.

**Figure S2.**
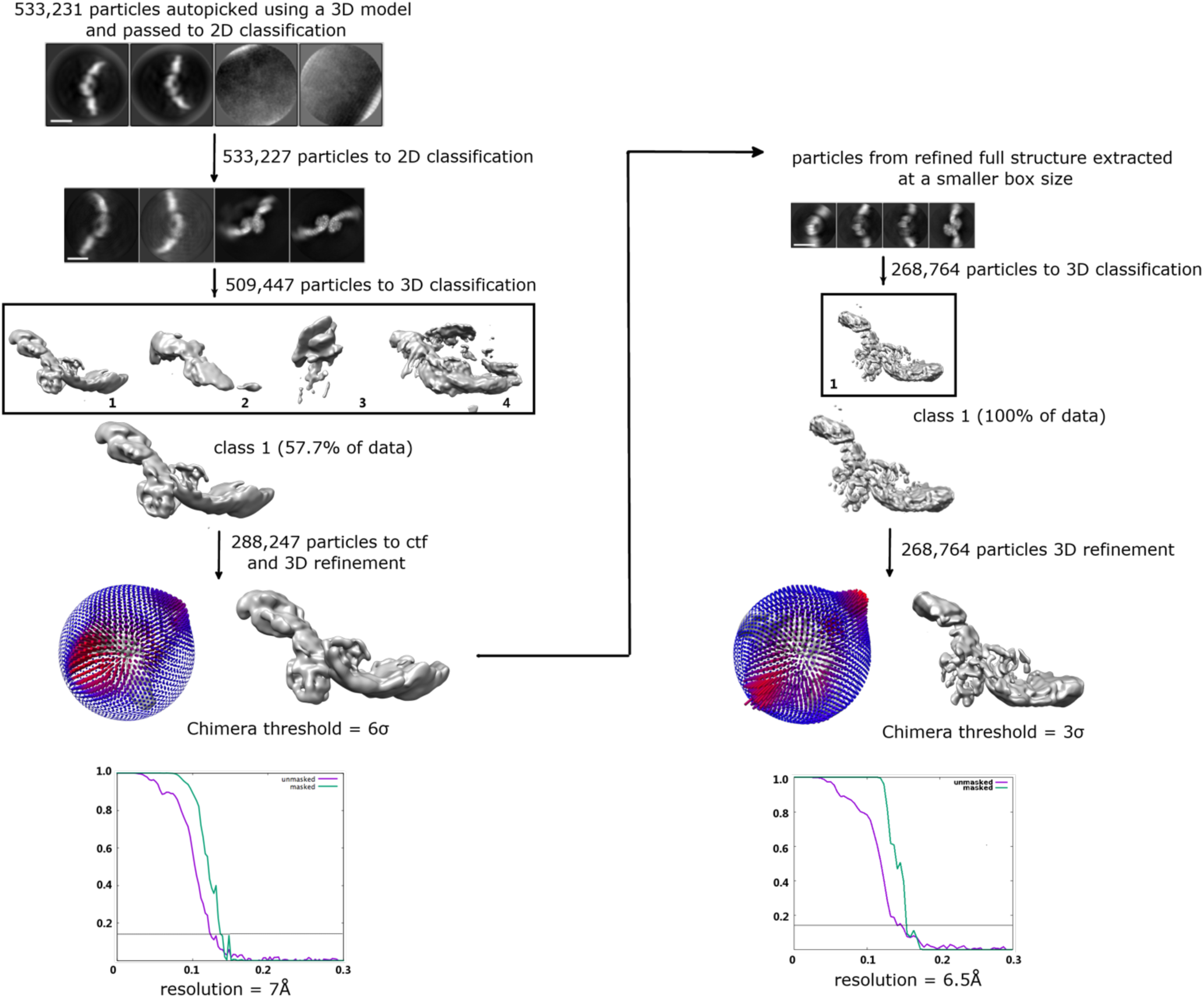
CryoEM image and data processing work flow for retromer dimers and sub-structure. Particles were auto-picked from a published dataset (Kendall, *Structure* 2020) using a 3D starting model of the retromer heterotrimer (details in Methods). Dimer particles were separated from other species during 2D classification, and 3D reconstructions were generated for both the intact dimer (left branch) and the VPS35/VPS35 sub-structure (right branch). Fourier Shell Correlation (FSC) plots showing masked and unmasked resolution estimates from RELION are shown for each structure or sub-structure; the grey line marks the 0.143 cut-off.

**Figure S3.**
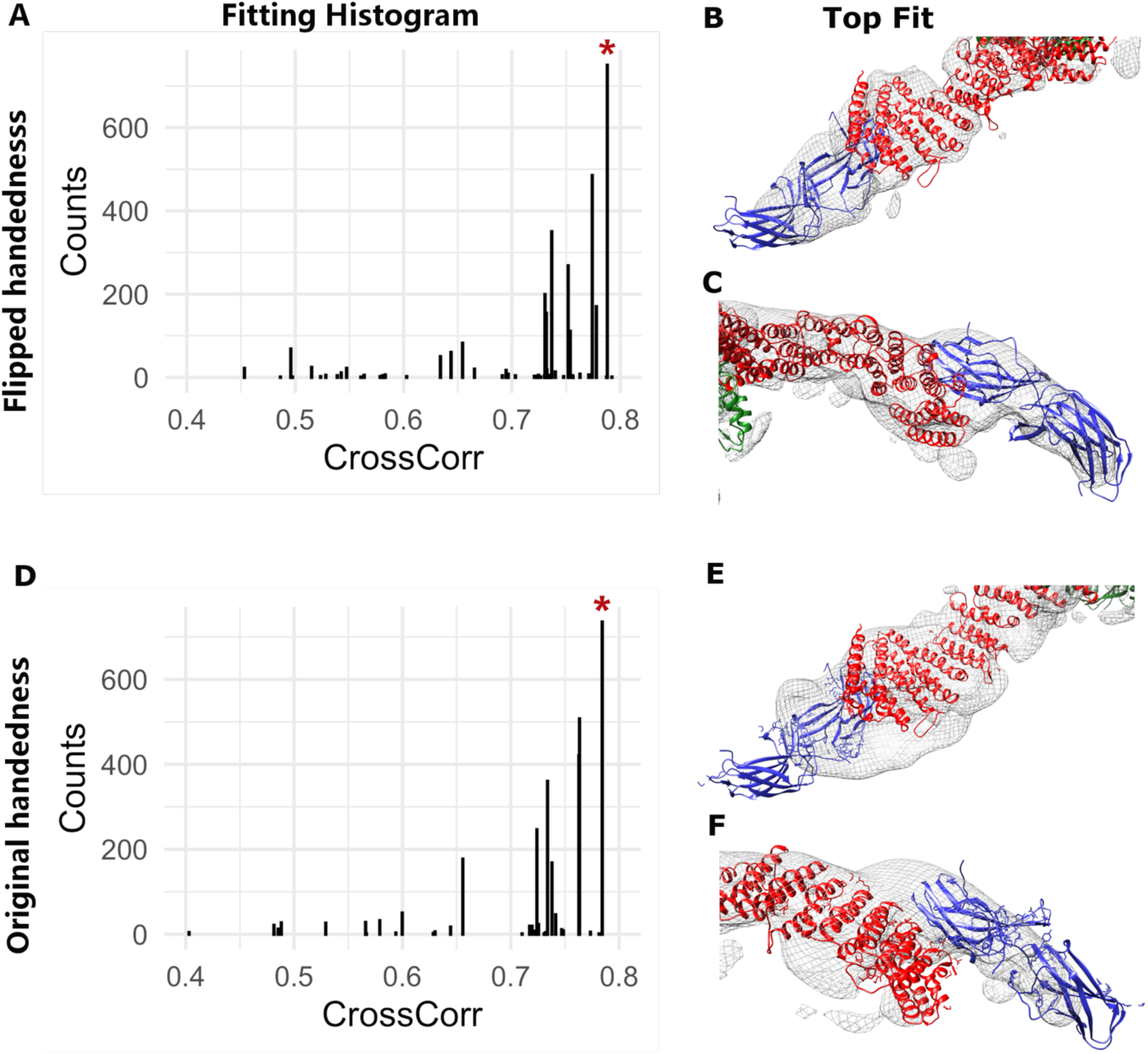
Analysis of retromer dimer map handedness. Map handedness was systematically analyzed by performing random rigid-body fits of real-space refined retromer models into each map (details in Methods). (A) Fitting histogram for flipped map handedness map showing counts of random rigid-body fits and cross-correlation values (Chimera). Magenta star marks top fit. (B,C) View of top fitted model in Coulomb potential map. Maps were generated using CCP4MG and are shown at 6σ contour level; VPS35 in red ribbons; VPS26 in blue ribbons; VPS29 in green ribbons. (D) Fitting histogram for original map handedness showing counts of random rigid-body fits and cross-correlation values (Chimera). (E, F) View of top fitted model in Coulomb potential map. Maps were generated using CCP4MG and are shown at 6σ contour level. This analysis suggests the flipped handedness (C, D) represents the correct handedness.

**Figure S4.**
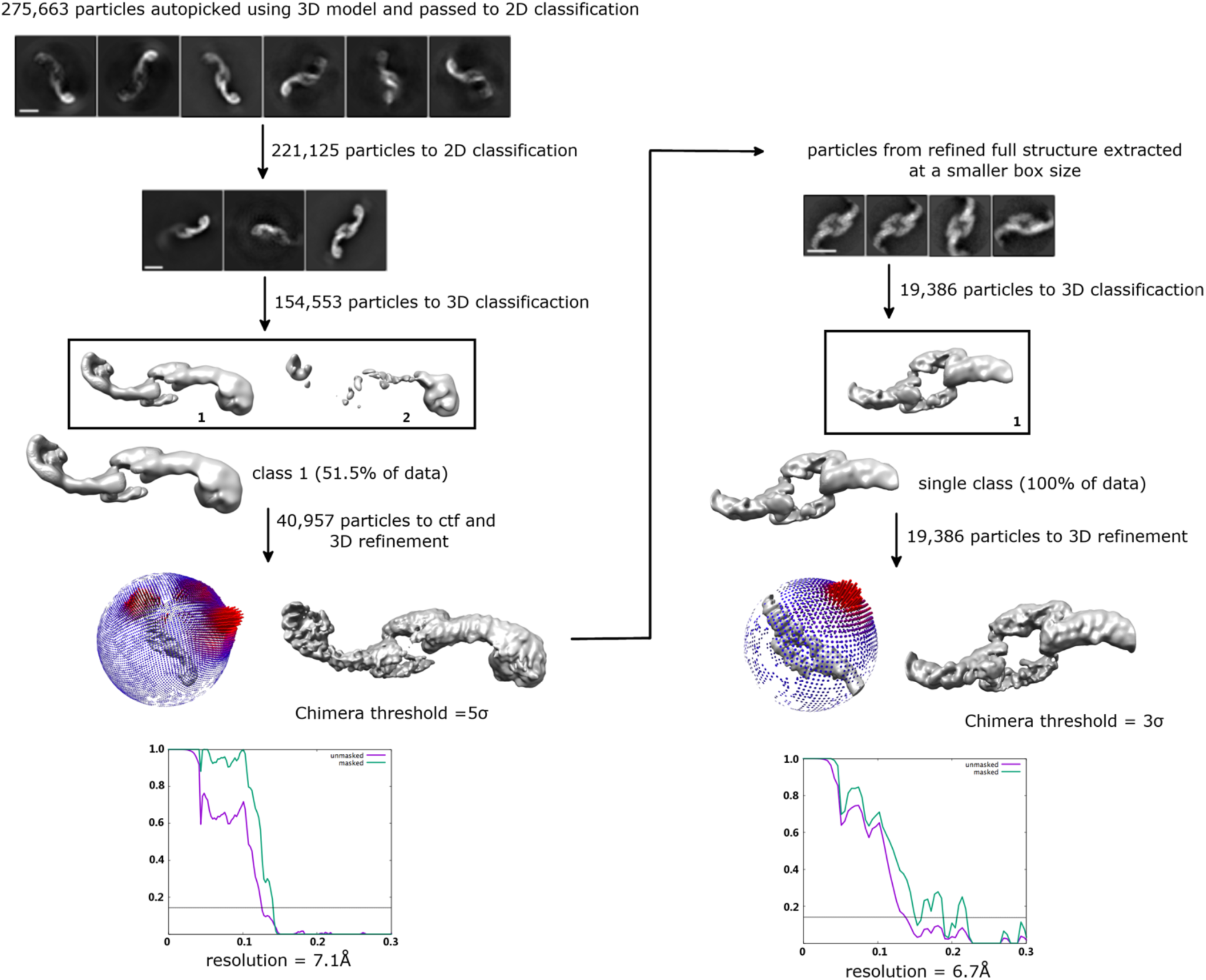
CryoEM image and data processing work flow for retromer 3KE mutant. Particles were auto-picked from a single dataset using a 3D starting model (details in Methods). 3KE mutant particles were separated from other biochemical species during 2D classification, and 3D reconstructions were generated for both the intact dimer (left branch) and the sub-structure focusing on the chain link (right branch). Fourier Shell Correlation (FSC) plots showing masked and unmasked resolution estimates from RELION are shown for each structure or sub-structure; the grey line marks the 0.143 cut-off.

**Figure S5.**
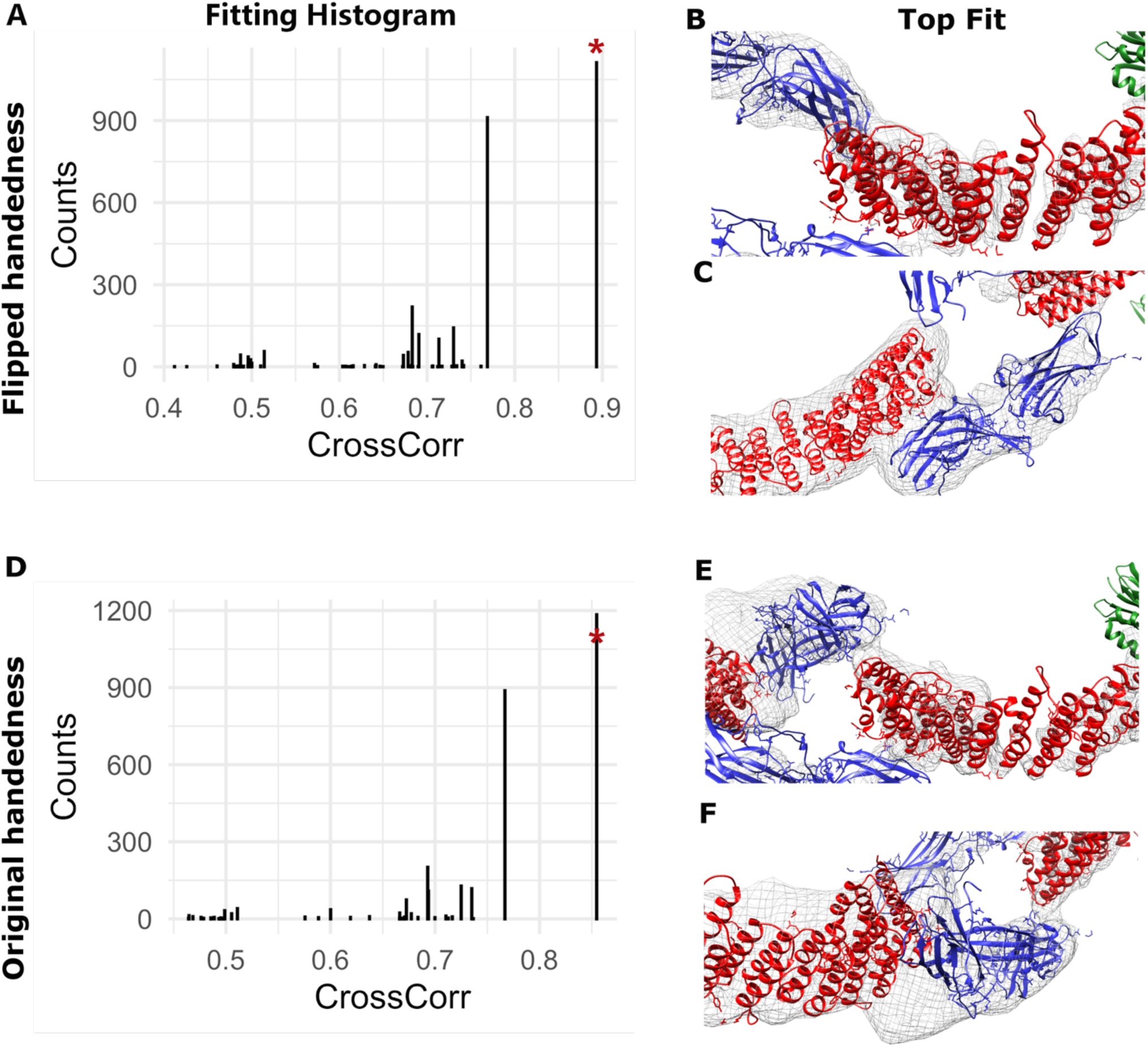
Analysis of retromer 3KE mutant map handedness. Map handedness was systematically analyzed by performing random rigid-body fits of real-space refined retromer models into each map (details in Methods); cross correlation coefficients and number of hits for each fit are shown. (A) Fitting histogram for flipped map handedness map showing counts of random rigid-body fits and cross-correlation values (Chimera). Magenta star marks top fit. (B, C) View of top fitted model in Coulomb potential map. Maps were generated using CCP4MG and are shown at 6σ contour level; VPS35 in red ribbons; VPS26 in blue ribbons; VPS29 in green ribbons. (D) Fitting histogram for original map handedness showing counts of random rigid-body fits and cross-correlation values (Chimera). (E, F) View of top fitted model in Coulomb potential map. Maps were generated using CCP4MG and are shown at 6σ contour level. This analysis suggests the flipped handedness (C, D) represents the correct handedness.

**Figure S6.**
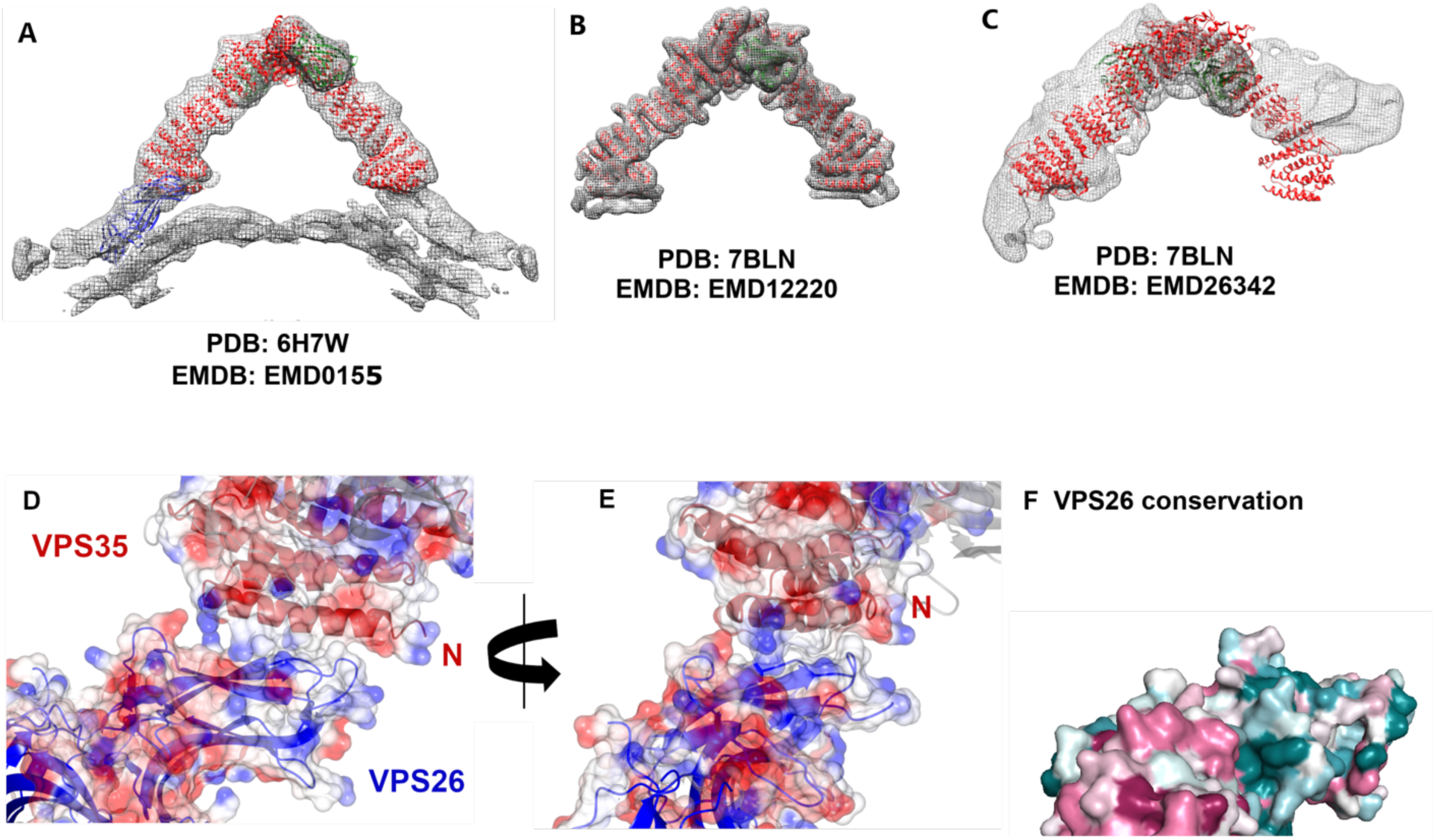
Analysis of retromer interfaces. (A, B) Equivalent views of published retromer arches are shown with their respective Coulomb potential maps from EMDB. (C) While individual retromer heterotrimers can be fitted as rigid bodies into the dimer map (cf. Figure 2; Figure S2), assembled arches are very poorly fitted as a single rigid body into the dimer map (rigid body fits in Chimera; full details in Discussion). (D, E) Two views rotated by 45 degrees of the N-VPS26A/N-VPS35 interface observed between a N-VPS35 (red ribbons) and N-VPS26A (blue ribbons) in a neighboring molecule; transparent electrostatic surface views are overlaid to demonstrate surfaces. The first helix of VPS35 (residues 12-36; labelled “N”) interacts primarily with two β-strands (residues 48-56; 105-111) and two loops (residues 56-63; 101-105) in N-VPS26A. The overall shape of each subunit is complementary to its binding partner. (F) VPS26 conservation (ConSurf) mapped onto the VPS26A structure. The N-terminal interface with VPS35 does not exhibit high conservation levels across eukaryotes, suggesting this interface may not occur in all organisms with retromer.

