## Supplementary material for "Improved mammalian retromer cryo-EM structures reveal a new assembly interface": Table 1

| | *3KE mutant and substructure* | *Dimer and VPS35/VPS35 substructure* | *Heterotrimer and VPS35/VPS35 substructure* | | |
|---|---|---|---|---|---|
| **Microscope** | ThermoFisher Titan Krios G3i Vanderbilt V-CEM | Titan Krios Krios2 NRAMM Data collection 1 | Titan Krios Krios2 NRAMM Data collection 1 | Titan Krios Krios3 NRAMM Data collection 2 | Titan Krios Krios3 NRAMM Data collection 3 |
| **Cs** | 2.7 mm | 2.7 mm | 2.7 mm | 2.7 mm | 2.7 mm |
| **Voltage** | 300 keV | 300 keV | 300 keV | 300 keV | 300 keV |
| **Detector** | ThermoFisher Falcon 3 direct electron detector | Gatan K2 Summit direct electron detector | Gatan K2 Summit direct electron detector | | |
| **Magnification** | 120,000x | 105,000x | 105,000x | 105,000x | 105,000x |
| **Pixel size** | 0.6811Å/pix | 1.096Å/pix | 1.096Å/pix | 1.06Å/pix (rescaled to 1.096Å/pix, motioncorr2 | 1.06Å/pix (rescaled to 1.096Å/pix, motioncorr2) |
| **Dose rate** | $1.4e^{-}/A^{2}$/sec | ~$8e^{-}/Å^{2}$/sec | ~$8e^{-}/Å^{2}$/sec | ~$8e^{-}/Å^{2}$/sec | ~$8e^{-}/Å^{2}$/sec |
| **Total dose** | $50e^{-}/A^{2}$ | $69.34e^{-}/Å^{2}$ | $69.34e^{-}/Å^{2}$ | $73.92e^{-}/Å^{2}$ | $73.92e^{-}/Å^{2}$ |
| **Tilt** | +/-30° | N/A | N/A | N/A | +/-15° |
| **Defocus range** | -0.8 to -2.6µm | -0.7 to 2.6µm | -0.7 to 2.6µm | -0.8 to -4.4µm | -0.8 to -4.7µm |
| **Number of micrographs** | 4,791 | 1,480 | 1,480 | 1,299 | 891 |
| **Total particles (autopicked)** | 275,633 | 533,231 | (250,500 particles selected from combined datasets; Kendall *et al*., 2020) | | 207,026 |
| **Box size (Å)** | 436x436 Å (3KE)<br>215x215 Å (substructure) | 351x351Å (dimer)<br>220x220 Å (substructure) | 241x241Å (heterotrimer)<br>180x180 Å (substructure) | | |
| **Particles in 2D classification** | 154,533 (3KE)<br>19,386 (substructure) | 509,447 (dimer)<br>288,247 (substructure) | 72,295 (heterotrimer)<br>69,381 (substructure) | | |
| **Particles in final 3D model** | 40,957 (3KE)<br>19,386 (substructure) | 288,247 (dimer)<br>268,764 (substructure) | 43,808 (heterotrimer)<br>69,381 (substructure) | | |
| **Symmetry** | C1 | C1 | C1 | | |
| **Map resolution (masked FSC 0.143, RELION)** | 7.1 Å (3KE)<br>6.7 Å (substructure) | 7.0 Å (dimer)<br>6.5 Å (substructure) | 4.9 Å (heterotrimer)<br>4.5 Å (substructure) | | |
| **B-factor (RELION)** | Not determined | Not determined | -114 (heterotrimer)<br>-226 (substructure) | | |
| **EMDB code** | EMD-26341 (3KE)<br>EMD-26340 (substructure) | EMD-26342 (dimer)<br>EMD-26343 (substructure) | EMD-24964 (heterotrimer)<br>EMD-26345 (substructure) | | |
| **PDB code** | N/A | N/A | XXXX | | |

**Table 1. Data collection and processing parameters.** Summary of data collection and processing parameters for all structures. The heterotrimer is presented here for completeness (Chen et al., *Sci Advances* 7, 2021).
