## Supplementary material for "Improved mammalian retromer cryo-EM structures reveal a new assembly interface": Table 2

| | | | | |
|---|---|---|---|---|
| All Atom Contacts | Clashscore, all atom: | 15.12 | | $49^{th}$ percentile* (N=1784, all resolutions) |
| | Clashscore is the number of serious steric overlaps (>0.4Å) per 1000 atoms. | | | |
| Protein Geometry | Poor rotamers | 31 | 2.81% | Goal: < 0.3% |
| | Favored rotamers | 1006 | 91.29% | Goal: > 98% |
| | Ramachandran outliers | 3 | 0.25% | Goal: < 0.05% |
| | Ramachandran favored | 1163 | 96.20% | Goal: > 98% |
| | Rama distribution Z-score | -2.35 ± 0.21 | | Goal: abs (Z score) < 2 |
| | MolProbity score^ | 2.28 | | $60^{th}$ percentile* (N=27675, 0Å - 99Å) |
| | Cβ deviations >0.25Å | 0 | 0.00% | Goal: 0 |
| | Bad bonds: | 0/10212 | 0.00% | Goal: 0% |
| | Bad angles: | 3/13740 | 0.02% | Goal: < 0.1% |
| Peptide Omegas | Cis Prolines: | 2/43 | 4.65% | Expected: ≤1 per chain, or ≤5% |
| Low-resolution Criteria | CaBLAM outliers | 15 | 1.3% | Goal: < 1.0% |
| | CA Geometry outliers | 3 | 0.25% | Goal: < 0.5% |
| Additional Validations | Chiral volume outliers | 0/1542 | | |
| | Waters with clashes | 4/100 | 4.00% | |

In the two column results, the left column gives the raw count, right column gives the percentage.

* $100^{th}$ percentile is the best among structures of comparable resolution; $0^{th}$ percentile is the worst. For clashscore the comparative set of structures was selected in 2004, for MolProbity score in 2006.

^ MolProbity score combines the clashscore, rotamer, and Ramachandran evaluations into a single score, normalized to be on the same scale as X-ray resolution.

**Table 2. Molprobity validation score summary for retromer heterotrimer structure.**
